# Arabidopsis 3-deoxy-D-*arabino*-heptulosonate 7-phosphate (DAHP) synthases of the shikimate pathway display both manganese- and cobalt-dependent activities

**DOI:** 10.1101/2024.10.06.616849

**Authors:** Ryo Yokoyama, Hiroshi A Maeda

## Abstract

The plant shikimate pathway directs a significant portion of photosynthetically assimilated carbon into the downstream biosynthetic pathways of aromatic amino acids (AAA) and aromatic natural products. 3-Deoxy-D-*arabino*-heptulosonate 7-phosphate synthase (DHS) catalyzes the first step of the shikimate pathway, playing a critical role in controlling the carbon flux from central carbon metabolism into the AAA biosynthesis. Previous biochemical studies suggested the presence of manganese- and cobalt-dependent DHS enzymes (DHS-Mn and DHS-Co, respectively) in various plant species. Unlike well-studied DHS-Mn, however, the identity of DHS-Co is still unknown. Here, we show that all three DHS isoforms of *Arabidopsis thaliana* exhibit both DHS-Mn and DHS-Co activities *in vitro*. A phylogenetic analysis of various DHS orthologs and related sequences showed that Arabidopsis 3-deoxy-D-*manno*-octulosonate-8-phosphate synthase (KDOPS) proteins were closely related to microbial Type I DHSs. Despite their sequence similarity, these Arabidopsis KDOPS proteins showed no DHS activity. Meanwhile, optimization of the DHS assay conditions led to the successful detection of DHS-Co activity from Arabidopsis DHS recombinant proteins. Compared to DHS-Mn, DHS-Co activity displayed the same redox dependency but distinct optimal pH and cofactor sensitivity. Our work provides biochemical evidence that the DHS isoforms of Arabidopsis possess DHS-Co activity.

## Introduction

The shikimate pathway connects central carbon metabolism to the biosynthesis of aromatic amino acids (AAAs, i.e. L-tyrosine, L-phenylalanine, and L-tryptophan) and AAA-derived natural products in plants. The shikimate pathway consists of seven enzymatic reactions to produce chorismate, the final pathway product and the last common precursor to all three AAAs (Herrmann, 1995; Tzin and Galili, 2010; Maeda and Dudareva, 2012; Yokoyama, 2024). The 3-deoxy-D-*arabino*-heptulosonate 7-phosphate synthase (DHS, also known as DAHP synthase) enzyme catalyzes the first committed step of the shikimate pathway. The condensation of phospho*enol*pyruvate (PEP) from glycolysis, and D-erythrose 4-phosphate (E4P) from the pentose phosphate pathways (mainly Calvin-Benson cycle in photosynthetic organisms) by the DHS enzyme generates DAHP, leading to the downstream six enzymatic steps into chorismate (Maeda and Dudareva, 2012; Yokoyama et al., 2022a). In plants, up to 30% of photosynthetically fixed carbon is directed into the shikimate pathway to support the high production of AAA-derived aromatic natural products, including lignin, an aromatic polymer that forms the secondary cell wall of plants (Razal et al., 1996; Vanholme et al., 2010). To enable flexible but strict regulation of the biosynthesis of AAAs and AAA-derived compounds, the DHS reaction plays a critical role as a gatekeeper to control carbon flux into the shikimate and the downstream AAA pathways (Yokoyama et al., 2022a). However, the regulation of plant DHS enzymes has only started to be investigated in depth in recent years.

Plant DHS activity has been investigated in various plant, dissecting its potential multi-layered complex regulatory systems. The main regulatory mechanisms controlling DHS activity in plants include negative feedback inhibition, redox regulation, and distinct cation cofactor sensitivity (Yokoyama et al., 2022a). In microbes, DHS enzymes are simply feedback inhibited by either AAA (Schoner and Herrmann, 1976; Paravicini et al., 1989; Bentley, 1990) On the other hand, plant DHS activity is subjected to a more complex negative feedback network mediated by several downstream aromatic compounds (Yokoyama et al., 2021; Yokoyama et al., 2022a). Plants utilize multiple DHS isoforms with different sensitivities to the downstream molecules likely to adjust carbon flux into the shikimate pathway (Yokoyama et al., 2021; El-Azaz et al., 2023). Identification of point mutations that attenuated the DHS negative feedback inhibition and thereby elevated AAA production in *Arabidopsis thaliana* revealed that DHS enzymes mediate the rate-limiting step in the shikimate and AAA biosynthetic pathways (Yokoyama et al., 2022b). In addition, plant DHS activity is subjected to redo regulation. Plant DHS activity often requires reducing power, such as dithiothreitol (DTT) and thioredoxin *f*, an isoform of the oxidoreductase proteins that reduce target proteins using redox power mainly derived from photosynthesis (Entus et al., 2002; Yokoyama et al., 2021). This redox dependency likely plays a regulatory role in activating DHS enzymes under photosynthesis-active conditions in the chloroplasts (Yokoyama et al., 2022a).

Another poorly characterized mechanism of DHS enzymes is their dual subcellular localization in cytosol and plastids. Enzymes in the shikimate and AAA pathways are dominantly localized in plastids (Rippert et al., 2009; Maeda and Dudareva, 2012; Yokoyama, 2024). On the other hand, many of the enzymes in the pathways, including DHS enzymes, have been additionally detected in the cytosol of various plant species to participate in cytosolic AAA metabolism (Lynch, 2022; Yokoyama et al., 2022a; Yokoyama, 2024). Previous studies separated chromatographic fractions of two distinct DHS activities requiring manganese (Mn) and cobalt (Co) ions as cofactors (Graziana and Boudet, 1980; Rubin and Jensen, 1985; Ganson et al., 1986; McCue and Conn, 1989; Doong et al., 1992; Doong and Jensen, 1992; Suzuki et al., 1996). These biochemical studies showed that Mn-dependent DHS (DHS-Mn) activity is highly enriched in plastidic fractions, whereas Co-dependent DHS (DHS-Co) activity is found in cytosolic fractions (Ganson et al., 1986). DHS-Mn activity of DHS recombinant proteins from Arabidopsis and some grass species has been well-characterized biochemically, demonstrating that DHS proteins participate in DHS-Mn activity *in vitro* (Yokoyama et al., 2021; El-Azaz et al., 2023). In contrast, enzymes responsible for DHS-Co activity remain unknown, with previous biochemical studies failing to detect DHS-Co activity in AtDHS recombinant proteins (Entus et al., 2002; Yokoyama et al., 2021).

Here, we investigated DHS-Co by first testing this activity in 3-deoxy-D-*manno*-octulosonate-8-phosphate synthase (KDOPS), a key enzyme mediating a key step in biosynthesizing one of the structurally divergent oligosaccharide side chains in cell wall rhamnogalacturonan II in bacteria and plants (Brabetz et al., 2000; Radaev et al., 2000; Raetz and Whitfield, 2002; Wu et al., 2004; Delmas et al., 2008). KDOPS proteins catalyze a reaction similar to DHS: the condensation of the phosphate sugar arabinose 5-phosphate (A5P) with PEP. Interestingly, previous publications revealed that the plant-derived fractions of DHS-Co, but not DHS-Mn, additionally exhibited the KDOPS activity (Doong et al., 1992; Doong and Jensen, 1992), leading us to the hypothesis that plant KDOPS enzymes may harbor DHS-Co activity. Our phylogenetic analysis showed that plant KDOPS proteins are evolutionarily close to microbial DHSs. Despite their sequence similarity, Arabidopsis KDOPS proteins exhibited neither DHS-Mn nor DHS-Co activities. In the meantime, recharacterization of AtDHS enzymes by optimizing assay conditions led to successful detection of DHS-Co activity. Furthermore, we found that the detected DHS-Co activity exhibit different biochemical properties from those of DHS-Mn, such as optimal pH and cofactor sensitivity, which probably made the detection of DHS-Co activity difficult in previous studies. These results demonstrate that DHS proteins exhibit DHS-Co activity, providing new biochemical insight into the identity of plant DHS-Co activity.

## Results

Earlier phylogenomic studies and structural comparison indicated that microbial DHS proteins are evolutionarily related to KDOPS proteins, which catalyze a similar condensation reaction to DHSs (Radaev et al., 2000; Wagner et al., 2000; Jensen et al., 2002). However, the information on their evolutionary relation with the plant DHS protein family remains limited. To elucidate the evolutional relation between DHSs and KDOPSs in plants and microbes, a cladogram tree of several plant DHS and KDOPS proteins as well as DHSs from microbes, including cyanobacteria, was constructed. Consistent with a previously characterized inter-kingdom DHS phylogenetic tree (Yokoyama et al., 2022a), *Escherichia coli* and *Saccharomyces cerevisiae* DHSs and cyanobacterial DHSs were divided into clusters of type Iα and type Iβ, respectively, whereas plant-type DHSs grouped into type II together with *Mycobacterium tuberculosis* DHS (**Figure 1**). Notably, plant KDOPS proteins were clustered next to the type Iβ DHS group together with bacterial KDOPSs (**Figure 1**). From this result, we hypothesized that plant KDOPS proteins may have DHS activity in addition to KDOPS activity.

**Figure 1.**
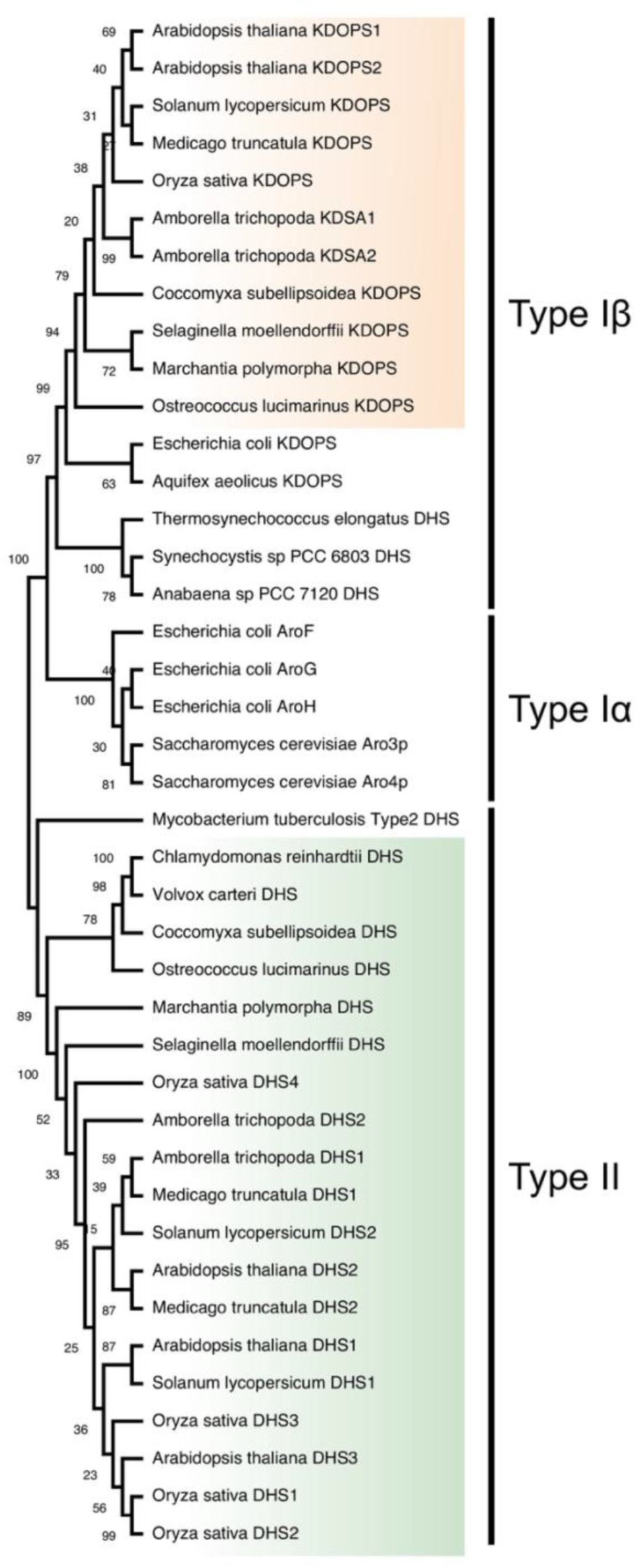
Plant KDOPs are closely related to Type Iβ microbial DHSs. A cladogram tree of DHS and KDOPS orthologs from plants and microbes was constructed using the Maximum likelihood method with 1,000 bootstrap replicates. DHS and KDOPS orthologs from eukaryotic photosynthetic organisms were highlighted in green and orange, respectively.

To test this hypothesis, recombinant proteins of two Arabidopsis KDOPS isoforms, KDOPS1 and KDOPS2 (AT1G79500 and AT1G16340, respectively), were expressed in *E. coli* and purified to examine their DHS activity *in vitro* (**Supplemental Figure 1**). KDOPS1 and KDOPS2 showed some DHS activities in the presence of Mn^2+^ or Co^2+^. However, these activities with Mn^2+^ or Co^2+^ were comparable with those in the presence of ethylenediaminetetraacetic acid (EDTA), a cation chelator, and more than 100-fold lower than AtDHS1 that was used as a positive control across different assay conditions of pH, redox, and cations (**Figure 2**). These findings indicate that, despite structural similarities between plant KDOSP and type I DHSs (**Figure 1**), Arabidopsis KDOPSs are unlikely to contribute to DHS-Co activity.

**Figure 2.**
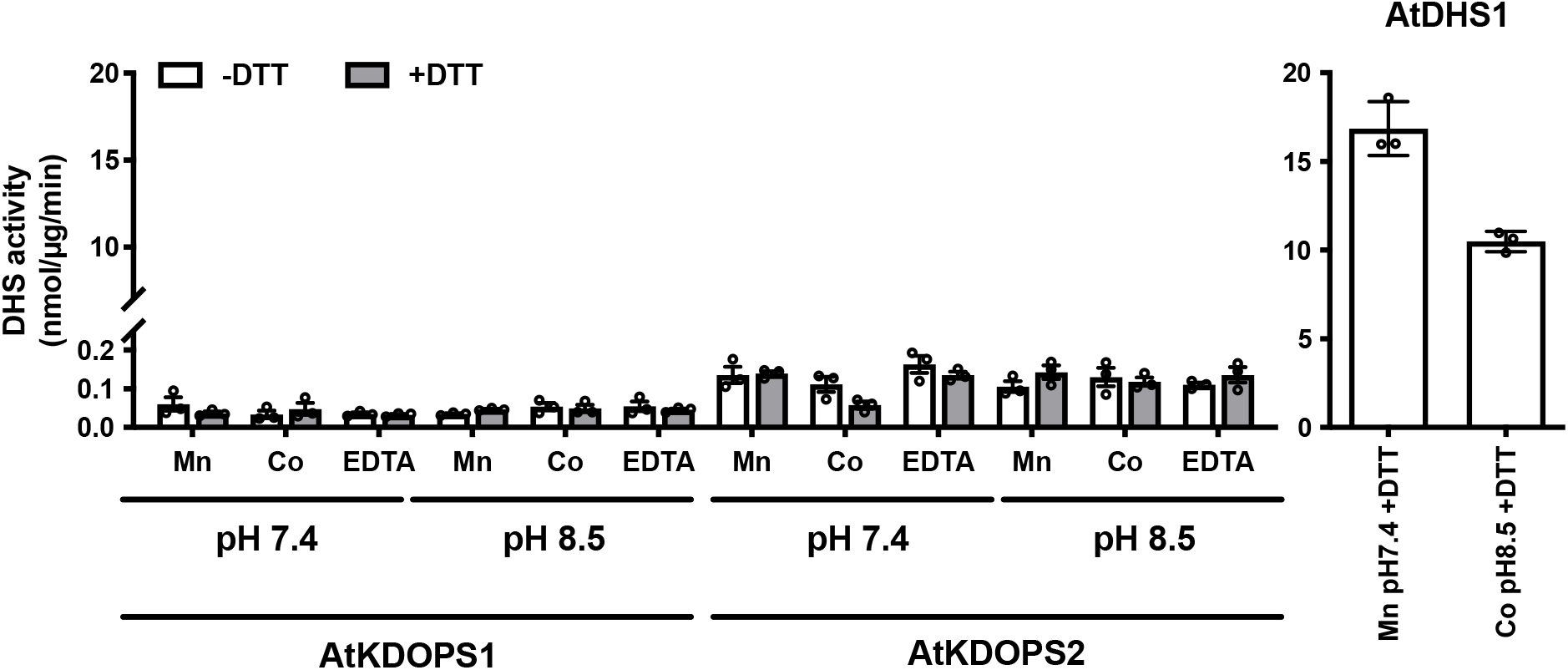
Arabidopsis KDOPS proteins lacked DHS-Mo or DHS-Co activities. Enzymatic assays of AtKDOPS1 and AtKDOPS2 using HEPES or EPPS buffers with Mn^2+^, Co^2+^ or EDTA in the absence or presence of DTT. DHS-Mn and DHS-Co activities from AtDHS1 were used as positive controls. Data are means ± SEM (*n* = 3 independent reactions). All the individual data points are shown as dots.

The lack of DHS-Co activity from the KDOPS recombinant enzymes led us to recharacterize the Arabidopsis DHS recombinant proteins using different types of buffers at distinct pH in the presence or absence of DTT. For this experiment, we first employed the colorimetric method to detect the production of DAHP as its oxidized form (Yokoyama et al., 2021). AtDHS1 and AtDHS2, two representatives of the three Arabidopsis DHS isoforms, showed the highest DHS activity with Mn^2+^ and DTT at pH 7.4 using 4-(2-hydroxyethyl)-1-piperazineethanesulfonic acid (HEPES) as a buffer reagent, whereas the residual DTT-independent activity was detected in AtDHS2 (**Figure 3A**). This result was consistent with the previous report (Yokoyama et al., 2021). With tris(hydroxymethyl)aminomethane (Tris)-based buffer at pH 8.5, none of the AtDHS proteins exhibited the DHS-Co activity (**Figure 3A**). Notably, when 4-(2-hydroxyethyl)-1-piperazinepropanesulfonic acid (EPPS) was used at pH 8.5, DHS-Co activity was observed in AtDHS1 and AtDHS2 in the presence of DTT, but not in the absence of DTT (**Figure 3A**).

**Figure 3.**
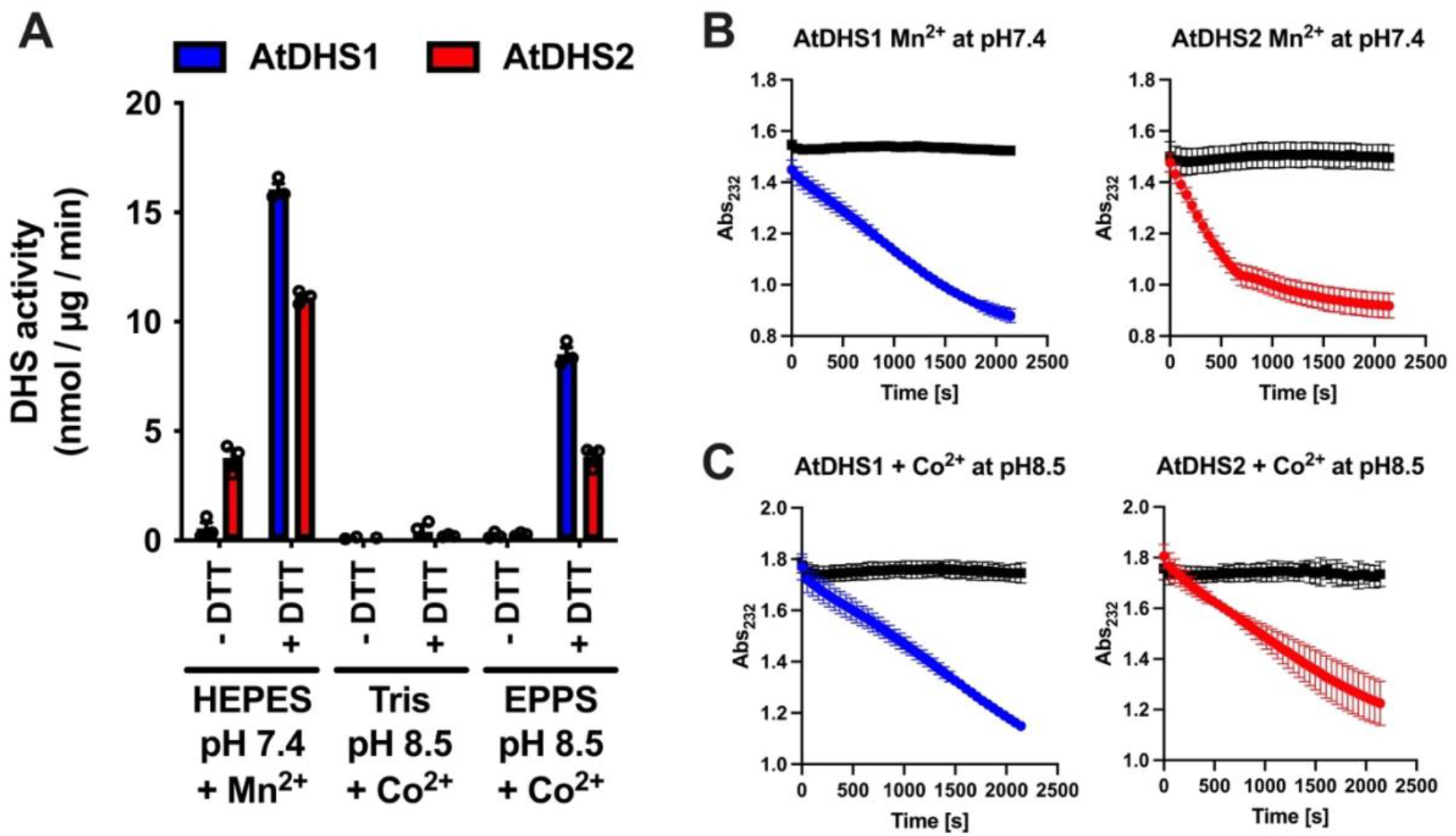
DHS-Co activity was detected at pH 8.5 in the presence of DTT. **(A)** Enzymatic assays of AtDHS1 and AtDHS2 using HEPES, Tris or EPPS buffers in the absence or presence of DTT. The colorimetric method was used to quantify DAHP production. **(B and C)** The PEP consumption from AtDHS1 (blue) and AtDHS2 (red) recombinant proteins was monitored at absorption 232 nm to validate (A) DHS-Mo and (B) DHS-Co activities at pH 7.4 and 8.5, respectively. Boiled inactive enzymes (black) were used as the negative controls for each reaction. Data are means ± SEM (*n* = 3 independent reactions). All the individual data points are shown as dots in (A).

To confirm this result using an independent mean, we carried out a different DHS assay method based on monitoring PEP consumption by reading the disappearance of the absorption at 232 nm (Schoner and Herrmann, 1976; El-Azaz et al., 2023). The PEP consumption was successfully monitored for AtDHS1 and AtDHS2 proteins in the presence of Mn^2+^ at pH 7.4 (**Figure 3B**), as previously performed (El-Azaz et al., 2023). Consistent with the result obtained by the colorimetric method (**Figure 3A**), DHS-Co activity from both AtDHS1 and AtDHS2 recombinant proteins, but not from their boiled inactive enzymes, was monitored as the PEP consumption at pH 8.5 (**Figure 3B**). These DHS enzymatic assays using two different methods firmly demonstrate that AtDHS1 and AtDHS2 isoforms display both DHS-Mn and DHS-Co activities *in vitro*.

Prior biochemical comparison between DHS-Mn and DHS-Co uncovered higher optimal pH and lower half maximal effective concentration (*EC*_50_) values of cation ions of DHS-Co than those of DHS-Mn (Ganson et al., 1986). To determine if these DHS-Co-specific biochemical features are also observed from AtDHS recombinant proteins, DHS assays were conducted using EPPS at different pHs from 7.6 to 8.7. All three AtDHS proteins exhibited the highest DHS-Co activity at 8.4 or 8.5 (**Figure 4**), which is consistent with previously reported higher optimal pH of DHS-Co isolated from several plants, including *Nicotiana silvestris, Spinacia oleracea, Vigna radiata*, and cultured carrot cells (Rubin and Jensen, 1985; Ganson et al., 1986; Doong et al., 1992; Suzuki et al., 1996). Saturation curves using the chloride salts of Mn^2+^ and Co^2+^ at different concentrations revealed that DHS-Co activity was stimulated more efficiently at lower Co^2+^ concentrations, with the *EC*_50_ values of 0.19, 0.038, and 1.10 μM for AtDHS1, AtDHS2, and AtDHS3, respectively. These *EC*_50_ values of DHS-Co were approximately 80-to 3000-times lower than those of DHS-Mn (15.44, 121, and 529 μM for AtDHS1, AtDHS2, and AtDHS3, respectively) (**Figure 5**). This trend in lower *EC*_50_ values of DHS-Co than DHS-Mn is in agreement with ones reported in *Nicotiana silvestris* (Ganson et al., 1986). Collectively, we successfully detected DHS-Co activity from AtDHS recombinant enzymes whose biochemical properties were similar to those activities previously detected from various plant tissue extracts.

**Figure 4.**
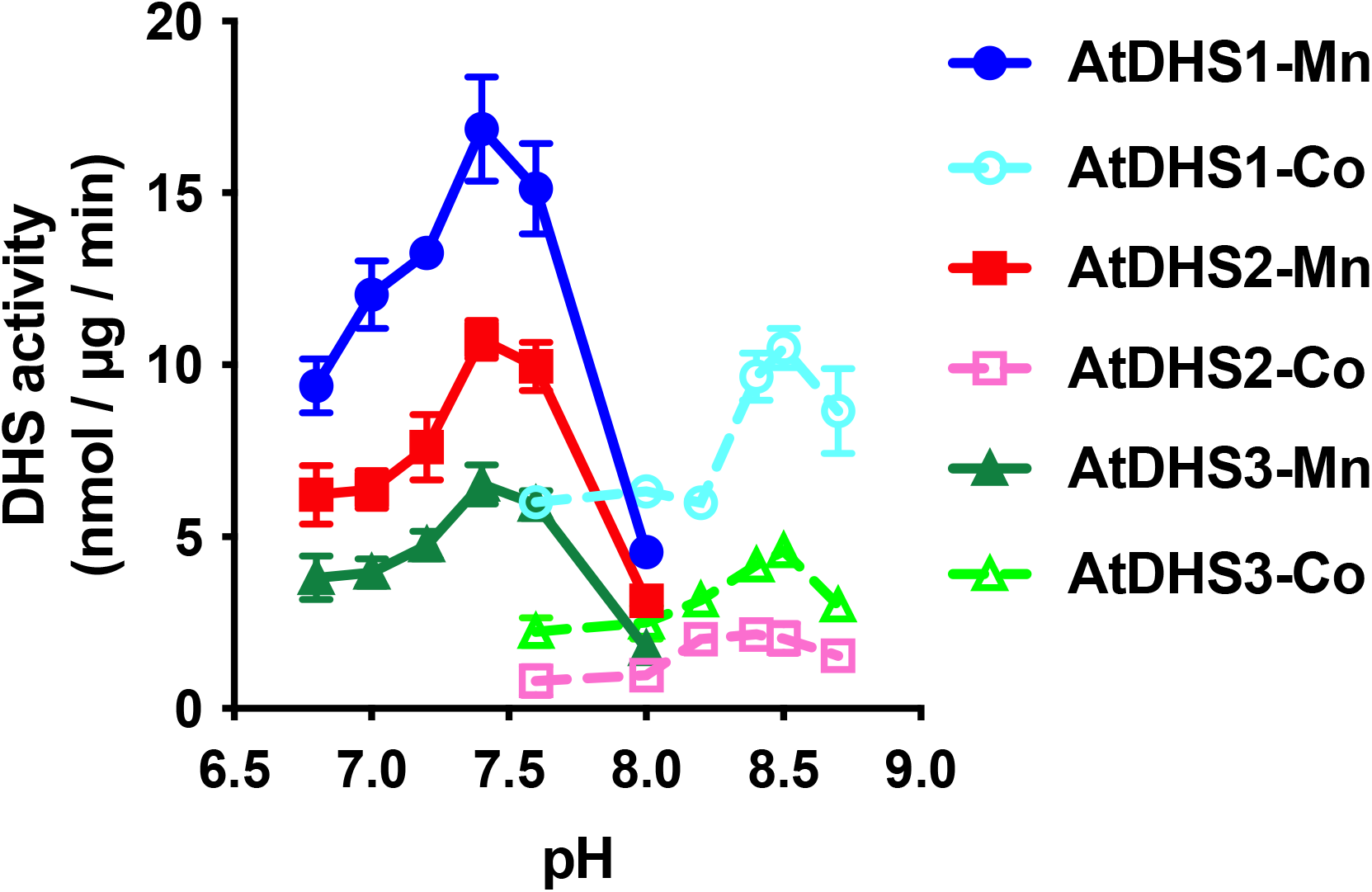
DHS-Co exhibited more alkaline optimal pH than DHS-Mn. Optimal pH of the recombinant Arabidopsis DHS enzymes in the presence of Mn^2+^ or Co^2+^. Enzymatic DHS-Mn and DHS-Co assays (solid and dashed lines, respectively) of Arabidopsis DHS1, DHS2, and DHS3 were conducted using HEPES buffer for pH 6.8 to 8.0 or EPPS buffer for pH 7.6 to 8.7, respectively. Data are means ± SEM (*n* = 3 independent reactions).

**Figure 5.**
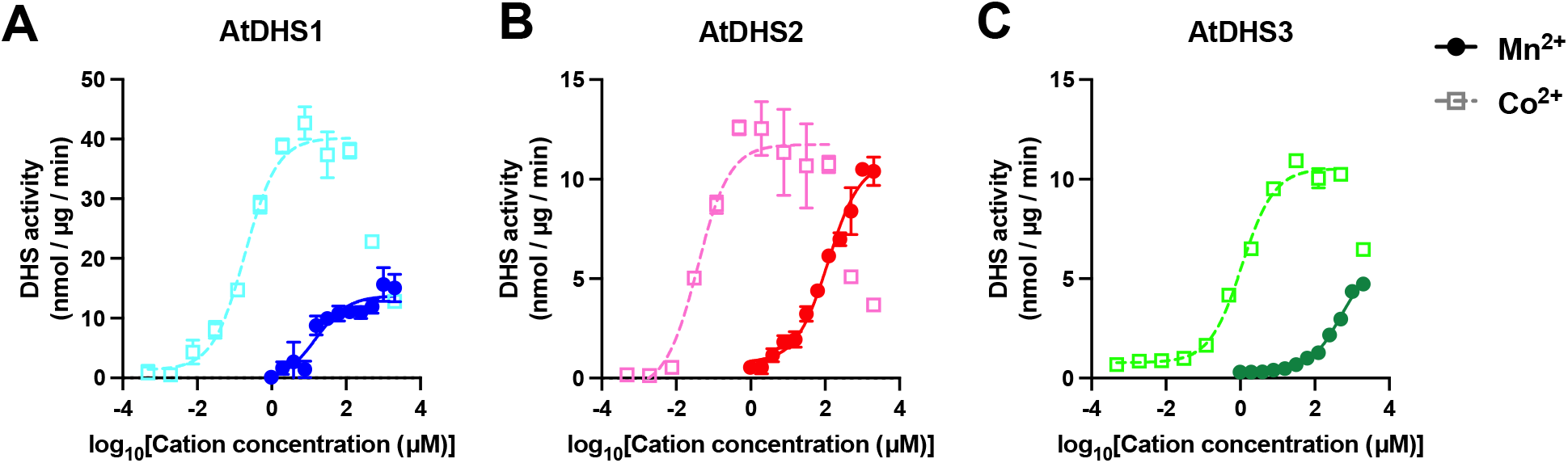
Co^2+^ ions stimulated DHS activity at much lower concentrations than Mn^2+^ ions. Enzymatic assays of AtDHS1 (A), AtDHS2 (B), and AtDHS3 (C) using a different concentration of Mn^2+^ or Co^2+^ (solid and dashed lines, respectively). The *EC*_50_ values DHS-Co were 0.19, 0.038, and 1.10 μM for AtDHS1, AtDHS2, and AtDHS3, respectively, while the *EC*_50_ values of DHS-Mn were 15.44, 121, and 529 μM for AtDHS1, AtDHS2, and AtDHS3, respectively). Data are means ± SEM (*n* = 3 independent reactions).

## Discussion

In this work, DHS-Co activity was successfully detected from AtDHS recombinant proteins using two different methods (**Figure 3**). Since DHS-Co has a higher optimal pH (pH 8.5) than DHS-Mn (pH 7.4) (**Figure 4**), it stands to reason that the prior studies failed to detect DHS-Co activity from the AtDHS recombinant proteins under conditions similar to the DHS-Mn assay. Higher optimal pH and lower *EC*_50_ of DHS-Co activity (**Figure 4 and 5**) are consistent with prior biochemical studies (Rubin and Jensen, 1985; Ganson et al., 1986; Doong et al., 1992; Suzuki et al., 1996), further supporting that DHS proteins are the component of and sufficient for DHS-Co activity. Furthermore, this study revealed that, like DHS-Mn activity, DTT was required for DHS-Co (**Figure 3A**). However, some prior studies reported that DHS-Co fractions derived from plant tissues did not require any redox regent for the DHS-Co assay (Rubin and Jensen, 1985; Ganson et al., 1986; Doong et al., 1992; Suzuki et al., 1996). The earlier biochemical work using DHS fractions isolated from *Nicotiana silvestris* discovered that DHS-Mn was more redox-demanding than DHS-Co (Ganson et al., 1986). This discrepancy may be due to the different redox states of protein fractions isolated from plant tissues and recombinant proteins isolated from bacteria. DHS proteins from plant tissues may already be sufficiently reduced in cells for DHS-Co, whereas the DHS recombinant proteins expressed in the bacterial host may not be sufficiently reduced to exhibit DHS-Co activity. How DHS-Co obtains redox energy in the cytosol, which is generally less reduced than in the chloroplast stroma (Wieloch, 2021), remains to be investigated.

The phylogenetic analysis suggests that DHS and KDOPS proteins share a distant evolutionary origin (**Figure 1**), consistent with their ability to mediate the similar condensation reactions of phosphate sugars with PEP (Radaev et al., 2000; Wagner et al., 2000). Because of their sequence similarity, a few bacterial KDOPS enzymes are able to react with other phosphate sugars, including E4P. While *E. coli* and *Aquifex aeolicus* KDOPS enzymes do not react with the other phosphate sugars (Ray, 1980; Duewel and Woodard, 2000), *Neisseria gonorrhoeae* KDOPS shows residual DHS activity (Subramaniam et al., 1998). Notably, a few previous studies reported that plant-derived DHS-Co fractions show KDOPS activity (Doong et al., 1992; Doong and Jensen, 1992). These previous biochemical reports of KDOPS and DHS proteins in bacteria and plants led to the hypothesis that plant DHS-Co activity might be derived from substrate ambiguity of plant KDOPS enzymes. However, KDOPS enzymes did not display a significant DHS activity under any of the pH, DTT, and cations conditions tested (**Figure 2**). These data together indicate that KDOPS enzymes are unlikely involved in DHS-Co activity. Given that plant and microbial DHS proteins form homo-dimers or tetramers (Suzuki et al., 1996; Jiao et al., 2020), DHS proteins may interact with KDOPS proteins in the cytosol due to their structural similarity and evolutionary relationship (**Figure 1**). This unfunctional but physical interaction between DHS and KDOPS proteins might result in co-elution of these two proteins and the detection of DHS-Co and KDOPS activities from the same protein fraction, as reported previously (Doong et al., 1992; Doong and Jensen, 1992).

While other proteins could still contribute to DHS-Co activity, this study provides biochemical evidence that Arabidopsis DHS recombinant proteins display both DHS-Mn and DHS-Co activities. DHS-Mn and DHS-Co fractions were separated from different plant tissues and species (Graziana and Boudet, 1980; Rubin and Jensen, 1985; Ganson et al., 1986; McCue and Conn, 1989; Doong et al., 1992; Doong and Jensen, 1992; Suzuki et al., 1996), implying that DHS-Mn or DHS-Co fractions may contain different protein components. Isolation of DHS-Mn and DHS-Co fractions from plant tissues and the following proteomics analysis will allow us to profile the components of DHS-Mn and DHS-Co in plants. Future investigations should also confirm the cytosolic DHS-Co localization and its metabolic impact on cytosolic AAA metabolism.

## Material and Method

### Preparation of protein expression vectors and recombinant proteins

Vectors for the expression of AtDHS1, AtDHS2, and AtDHS3 recombinant proteins are the same as what we used previously (Yokoyama et al., 2021). For the expression of His-tagged AtKDOPS1 and AtKDOPS2 recombinant proteins in *E. coli*, the full-length CDS fragments were gene-synthesized and cloned into pET100/D-TOPO (Thermo Fisher Scientific, Waltham, MA, USA). The expression and purification of the recombinant proteins were conducted as described previously (Yokoyama et al., 2021).

### Enzymatic assays

Unless otherwise noted, DHS enzymatic activity was monitored by the colorimetric method as previously described, (Yokoyama et al., 2021) with some modifications. The enzyme solution (7.7 μL) containing 0.01-0.05 μg proteins was used for the assays. After adding 0.5 μL of 0.1 M DTT or the same volume of H_2_O, the samples were further incubated at room temperature for 15 min. During these incubations, the substrate solution containing 50 mM HEPES pH 7.4, 2 mM MnCl_2_, 4 mM E4P, and 4 mM PEP for DHS-Mn or 50 mM EPPS pH 8.5, 2 mM CoCl_2_, 4 mM E4P, and 4 mM PEP at final concentration was preheated at 37°C. The enzyme reaction was started by adding 6.8 μL of the substrate solution (total 15 μL reaction volume), then incubated at 37°C for 20 min, and terminated by adding 30 μL of 0.6 M trichloroacetic acid. After a brief centrifugation, 5 μL of 200 mM NaIO_4_ (sodium meta-periodate) in 9 N H_3_PO_4_ was added to oxidize the enzymatic product and incubate at 25°C for 20 min. To terminate the oxidation reaction, 20 μL of 0.75 M NaAsO_2_ (sodium arsenite), which was dissolved in 0.5 M Na_2_SO_4_ and 0.05 M H_2_SO_4_, were immediately mixed. After 5 min of incubation at room temperature, one-third of the sample solution was transferred to a new tube to be mixed with 50 μL of 40 mM thiobarbituric acid and incubated at 99°C for 15 min in a thermal cycler. The developed pink chromophore was extracted by adding 600 μL of cyclohexanone in eight-strip solvent-resistant plastic tubes, mixing vigorously and centrifuging them at 4,500 × *g* for 3 min to separate the water- and cyclohexanone-fractions. The absorbance of the pink supernatant was read at 549 nm in a microplate reader (Infinite 200 PRO, TECAN, Männedorf, Switzerland). DAHP production was calculated based on its molar extinction coefficient at 549 nm (ε = 549 nm) of 4.5 × 104/M/cm. Reaction mixtures with boiled enzymes were run in parallel and used as negative controls to estimate the background signal for recombinant enzymes.

DHS enzymatic activity was also measured by the PEP-consumption method as previously described (Schoner and Herrmann, 1976; El-Azaz et al., 2023), with some modifications. The enzyme solution (33.5 μL) containing 50 mM HEPES pH 7.4 and 2 mM MnCl_2_ or 50 mM EPPS pH 8.5 and 2 mM CoCl_2_ (concentrations refer to the final reaction volume of 50 μL) and excess amounts of AtDHS recombinant proteins (0.5 μg) were prepared in a 96-well microplate, and 1.5 μL of 100 mM DTT stock were added and incubated at room temperature for 5 minutes to reduce AtDHS proteins. After adding 5 μL of 15 mM PEP solution, the plate was incubated in the plate reader at 37°C for 5 min, followed by the absorption measurement at 232 nm. The reaction was initiated by adding 5 μL of 20 mM E4P solution, and the absorption was recorded at 37°C at every-30-second intervals with shaking in between. Boiled inactive enzymes were used as the negative controls for each reaction.

### Construction of DHS and KDOPS cladogram tree

KDOPS orthologs were first identified by BlastP searches utilizing the amino acid sequence of AtKDOPS1 as a query against Phytozome (Goodstein et al., 2012). DHS ortholog sequences were used from a previous study (Yokoyama et al., 2022a). All the obtained sequences were then used to construct a tree of DHS and KDOPS proteins using MEGA X (Kumar et al., 2018) and are available as a FASTA file in Supplemental Data Set S1. The sequences were aligned by the MUSCLE algorithm and then constructed into the tree based on the maximum-likelihood method with 1,000 bootstrap replicates.

### Accession numbers

Sequence data from this article can be found in the EMBL/GenBank data libraries under the following accession numbers: AtDHS1 (AT4G39980), AtDHS2 (AT4G33510), AtDHS3 (AT1G22410), KDOPS1 (AT1G79500), and KDOPS2 (AT1G16340).

## Acknowledgments

The authors are grateful to Dr. Jorge El-Azaz (University of Wisconsin-Madison) for his critical reading.

## Contribution

R.Y. and H.A.M. designed the research. R.Y. performed the experiments. R.Y. analyzed the data, and R.Y. and H.A.M. wrote the manuscript.

Conflict of interest statement. None declared.

