## Supplemental Figure 1 for "Arabidopsis 3-deoxy-D-*arabino*-heptulosonate 7-phosphate (DAHP) synthases of the shikimate pathway display both manganese- and cobalt-dependent activities"

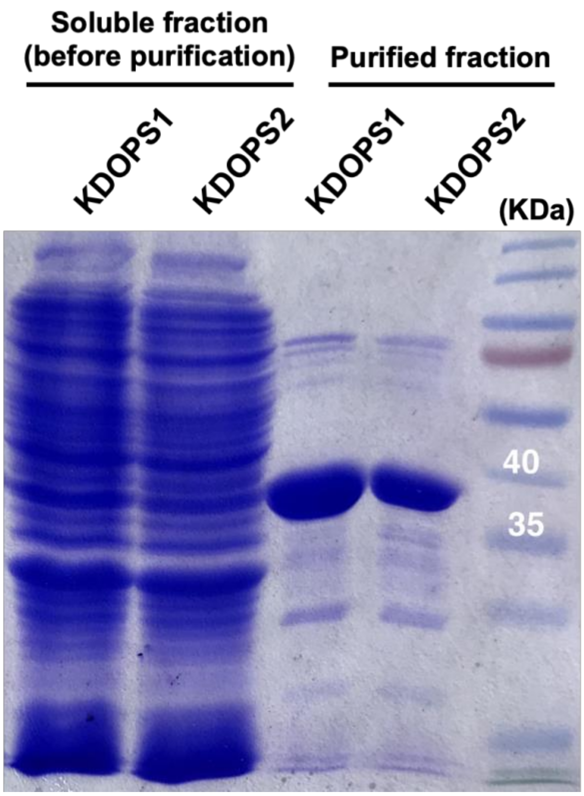


**Supplemental Figure 1. A gel image of purified Arabidopsis KDOPS1 and KDOPS2 recombinant proteins.**

The SDS-PAGE gel image of soluble fractions (left two samples) of bacteria expressing KDOPS1 and KDOPS2 recombinant proteins before being subjected to His-tag affinity purification and purified KDOPS1 and KDOPS2 recombinant proteins (right two samples). The expected protein sizes of His-tagged KDOPS1 and KDOPS2 recombinant proteins are 35.7 and 35.8 kDa, respectively.
